# A single cell transcriptional roadmap for human pacemaker cell differentiation

**DOI:** 10.1101/2021.12.28.474383

**Authors:** Alexandra Wiesinger, Jiuru Li, Lianne Fokkert, Priscilla Bakker, Arie O. Verkerk, Vincent M. Christoffels, Gerard J.J. Boink, Harsha D. Devalla

## Abstract

Each heartbeat is triggered by the sinoatrial node, the natural pacemaker of the heart. Animal models have revealed that pacemaker cells share a common progenitor with the (pro)epicardium, and that the pacemaker cardiomyocytes further diversify into “transitional”, “tail” and “head” subtypes. However, the underlying molecular mechanisms are poorly understood. Here, we studied the differentiation of human induced pluripotent stem cells into pacemaker cardiomyocytes. Single cell RNA sequencing identified the presence of myocardial populations resembling subtypes present in the formed sinoatrial node, and in addition revealed a side population of (pro)epicardial cells. Time-course trajectory analysis uncovered a role for WNT signaling in determining myocardial versus proepicardial cell fate. We experimentally demonstrate that presence of WNT signaling prior to the branching point of a common progenitor enhances proepicardial cell differentiation at the expense of myocardial pacemaker cells. Furthermore, we uncover a role for TGFβ and WNT signaling in differentiation towards transitional and head pacemaker subtypes, respectively. Our findings provide new biological insights into human pacemaker differentiation, open avenues for complex disease modeling and inform regenerative approaches.

## Introduction

The human heart beats about three billion times in an average life span. Each heartbeat is triggered by the electrical impulses generated by the sinoatrial node (SAN), referred to as the primary pacemaker of the heart. Dysfunction of the SAN results in potentially life-threatening cardiac arrhythmias (Choudhury *et al*, 2015) and current treatment with the implantation of electronic pacemakers is suboptimal (Cingolani *et al*, 2018). A better understanding of the origin, composition and function of the human SAN will enable the development of effective therapies. Previous studies have revealed that the SAN is a complex heterogeneous structure composed of both myocardial and non-myocardial cells such as fibroblasts, smooth muscle cells etc., which contribute to its function (Wiese *et al*, 2009; Bressan *et al*, 2018; Goodyer *et al*, 2019). The mechanisms that regulate the development of the various cell types of the SAN niche remain largely unknown.

The pacemaker cells of the SAN originate from a *Tbx18*^+^ progenitor population that also gives rise to proepicardial cells (van Wijk *et al*, 2009; Mommersteeg *et al*, 2010). Moreover, proepicardium-derived mesenchymal cells have been shown to be integral for remodeling and sustained electrical activity of the SAN (Bressan *et al*, 2018). In chicken development, bone morphogenetic protein (BMP) and fibroblast growth factor (FGF) signaling have been shown to orchestrate the separation of myocardial and proepicardial cells (Kruithof *et al*, 2006; van Wijk *et al*, 2009). *In vitro* studies using human pluripotent stem cells point to a crosstalk between BMP, retinoic acid (RA) and wingless-related integration site (WNT) signaling in the differentiation of pacemaker and proepicardial cells (Wiesinger *et al*, 2021).

Within the cardiomyocyte fraction of the SAN, there are distinct subpopulations such as head, tail and transitional cells (Komosa *et al*, 2021). The pacemaker cells in the SAN-head population express the T-box transcription factors *Tbx18* and *Tbx3.* This region is also distinct from all other cardiomyocytes in the heart as it lacks the expression of *Nkx2-5* (Wiese *et al*, 2009). The SAN-tail located inferior to the SAN-head expresses *Tbx3* and *Nkx2-5* but is devoid of *Tbx18* (Wiese *et al*, 2009; Goodyer *et al*, 2019). Furthermore, transitional cells (SAN-TZ) with transcriptional and functional properties intermediate to that of pacemaker cells and the adjacent atrial myocardium have also been reported, which are believed to play a critical role in transmitting the electrical impulses from the SAN to the adjacent atrial myocardium (Boyett *et al*, 2000; Csepe *et al*, 2016; Goodyer *et al*, 2019; Li *et al*, 2019).

Whilst the shared origin of pacemaker cells with proepicardium and further differentiation of the pacemaker cells to distinct subpopulations is recognized, the mechanisms underlying these processes are poorly understood. Moreover, the vast majority of data regarding the development of the SAN is derived from animal models (van Eif *et al*, 2018) and our insights into human SAN development are very limited (Csepe *et al*, 2016; Chandler *et al*, 2009; Sizarov *et al*, 2011; van Eif *et al*, 2019). Differentiating human induced pluripotent stem cells (hiPSCs) are an excellent model to study human heart development *in vitro* providing easy access to early developmental stages and allowing the reconstruction of cell fate decisions.

Here, we show that the differentiation of hiPSCs to SAN cardiomyocytes (SANCM) recapitulates developmental programs with remarkable fidelity. Single cell RNA sequencing (scRNA-seq) demonstrated that the differentiated cell pool contains myocardial populations resembling pacemaker cell types in the different subdomains of the *in vivo* SAN, i.e., SAN-head, SAN-tail and SAN-TZ cells, in addition to a non-myocardial side population of pro-epicardial cells, reflecting their shared ontogeny. Using trajectory inference analysis tool URD, we provide a transcriptional roadmap of these cell types and identify that the fate decision of a common progenitor towards myocardial or proepicardial lineages is determined by WNT signaling. Importantly, the branching of the myocardial group to various SANCM subpopulations also appeared to follow a temporal order influenced at least in part by TGFβ and WNT signaling.

Our results provide insight into the specification and diversification of human pacemaker cells. The ability to obtain the various sub populations and steer this differentiation process offers opportunities for assembling advanced *in vitro* models to better understand SAN function in health and disease and will further strengthen the basic framework for the development of regenerative therapies.

## Results

### Differentiation of hiPSCs to sinoatrial nodal and ventricular cardiomyocytes

Differentiation of hiPSCs towards *MESP1*^+^ mesoderm was initiated by activating Activin/Nodal, BMP and WNT signaling, as previously described (Devalla *et al*, 2015, 2016). To steer mesoderm towards a cardiomyocyte fate, WNT signaling was inhibited using XAV 939 for 96 hours, which resulted in predominantly ventricular-like cardiomyocytes (VCM). To direct mesoderm towards SANCM, we treated cultures with BMP4, RA, WNT inhibitor (XAV 939), FGF inhibitor (PD173074), and ALK5 inhibitor (SB431542) for 48 hours from day 4 to day 6 (Fig. 1A) (Protze *et al*, 2017). Contracting cardiomyocytes were observed from day 10 onwards and phenotypical differences in beating rates were apparent; SANCM monolayers exhibited faster beating rates in contrast to slower beating rates of VCM monolayers. TNNT2 expression was used as a measure of differentiation efficiency and flow cytometry analysis on day 19 demonstrated the presence of 60 – 90% cardiomyocytes in both groups (Fig. 1B).

**Figure 1.**
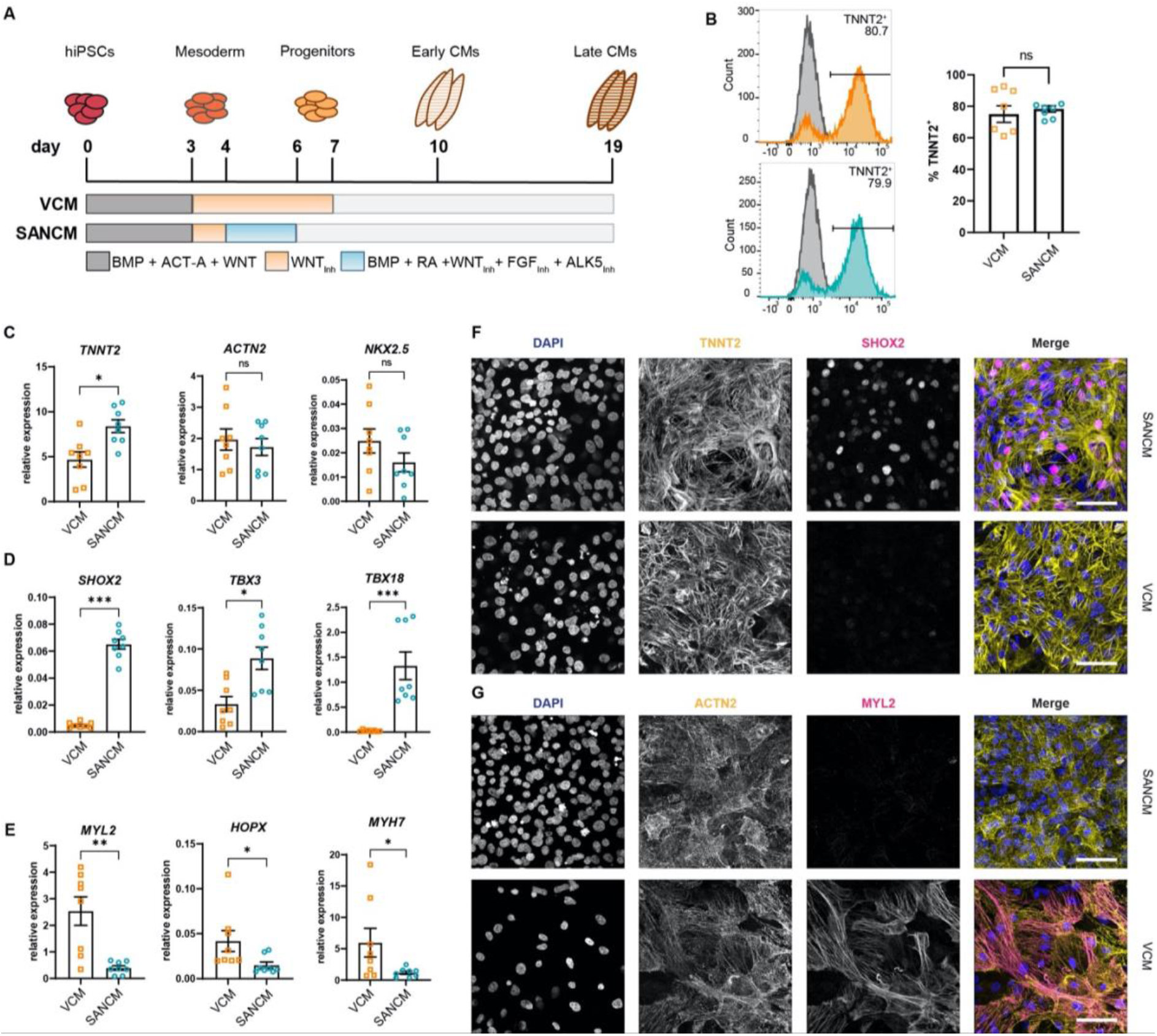
Differentiation of human induced pluripotent stem cells (hiPSCs) to sinoatrial node (SANCM) and ventricular cardiomyocytes (VCM). (A) Schematic representation of protocols used to differentiate hiPSCs to VCM and SANCM. (B) Representative histograms (left) and summarized data (right) showing percentage of TNNT2^+^ cells in VCM (orange) and SANCM (blue) at day 19 of differentiation. A corresponding IgG isotype antibody was used as negative control for flow cytometry (grey). n = 7 independent differentiations. Error bars, s.e.m. Mann-Whitney U test: P > 0.05 (ns). (C-E) RT-qPCR depicting expression of pan cardiomyocyte genes (C), SAN-associated genes (D) and ventricular-associated genes (E) at day 19 of differentiation. n = 8 independent differentiations; corrected to GEOMEAN of reference genes RPLP0 and GUSB. Error bars, s.e.m. Mann-Whitney U test: P < 0.05 (*), P < 0.005 (**), P < 0.0005 (***). (F-G) Immunofluorescence stainings demonstrating the expression of nuclear stain DAPI, SHOX2 and TNNT2 (F), MYL2 and ACTN2 (G), in SANCM and VCM. Scale bars, 50 μm.

To assess cardiomyocyte (CM) identity, gene expression profiling was performed by RT-qPCR. Both CM subtypes expressed sarcomeric genes *TNNT2*, *ACTN2* and the transcription factor *NKX2-5* (Fig. 1C). Although *NKX2-5* expression was generally lower in SANCM compared with VCM, the difference was not statistically significant. The expression of transcription factors *SHOX2*, *TBX3*, *TBX18* and *ISL1*, each required for proper SAN function (van Eif *et al*, 2018), was significantly higher in SANCM, indicating a SAN-like phenotype (Fig. 1D, S1A). VCM identity was verified by the expression of genes enriched in the ventricles, such as *MYL2*, *HOPX* and *MYH7* (Fig. 1E). In line with the above findings, immunofluorescence staining confirmed that SHOX2 and ISL1 are predominantly expressed in SANCM, whereas MYL2 expression was exclusively found in VCM (Fig. 1F and 1G and S1B).

### SANCM and VCM display distinct electrophysiological properties

Besides transcription factors, a number of ion channel genes are differentially expressed between the SAN and the ventricles, which confer distinct electrophysiological properties. The expression of *HCN1* and *HCN4*, which contribute to cardiac funny current I_*f*_, implicated in pacemaking, was significantly higher in SANCM compared with VCM. Similarly, the L-type and T-type Ca^2+^ channel genes *CACNA1D* and *CACNA1G*, respectively, as well as the inward rectifying K^+^ channel Kir3.1, encoded by *KCNJ3*, were significantly upregulated in SANCM compared with VCM (Fig. 2A). On the contrary, expression of *SCN5A*, the gene encoding cardiac Na^+^ channel Na_V_1.5, was higher in VCM (Fig. 2A). Consistently, action potential parameters (analyzed as in Fig. 2B) of SANCM and VCM measured by single cell patch-clamp confirmed expected subtype-specific electrophysiological differences. Representative traces of spontaneous action potentials are shown in Fig. 2C, demonstrating shorter cycle length in SANCM (496.6 ± 33.0 ms, mean ± SEM, n=12) compared with VCM (1241.5 ± 111.7 ms, n=12) (Fig. 2D). Consistent with a SAN phenotype, the maximum diastolic potential (MDP) was less negative in SANCM (−62.5 ±1.9 mV) compared with VCM (−69.9 ±1.4 mV). Furthermore, SANCM displayed a lower action potential amplitude (APA) and slower upstroke velocity (Vmax; 5.2 ± 0.9 V/s SANCM vs 23.1 ± 3.7 V/s VCM). Notably, MDPs and Vmax recorded in SANCM are similar to freshly isolated human SAN cells (Verkerk *et al*, 2007). On the contrary, longer action potential durations at 20, 50 and 90% repolarization (APD20, APD50, and APD90 respectively) characterized the VCM (Fig. 2D, Table S1). In addition, treatment with 3 μM ivabradine (IVA), an I_*f*_ channel blocker (Bucchi *et al*, 2002), resulted in a significant increase in cycle length in SANCM (baseline [BL]: 491.1 ± 76.8 ms; IVA: 771.9 ± 124.7 ms, n = 6), whereas cycle length in VCM was unaffected (BL: 838.0 ± 110.9 ms; IVA: 817.9 ± 114.0 ms) (Fig. 2E and Table S2). Taken together, these results affirm the cellular identities expected for SANCM and VCM.

**Figure 2.**
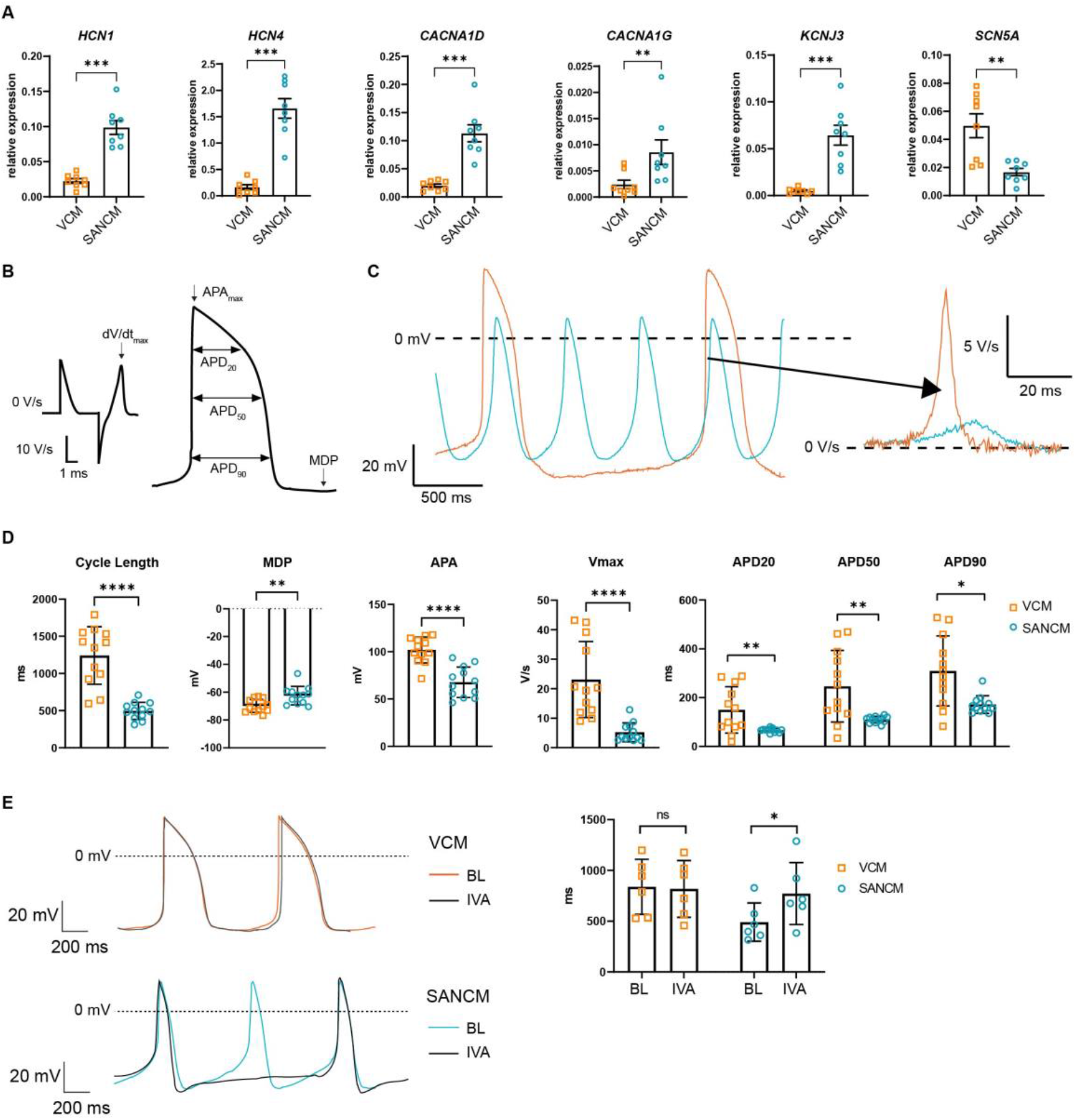
Electrophysiological characterization of SANCM and VCM. (A) RT-qPCR showing expression of ion channel genes at day 19 of differentiation. N = 8 independent differentiations; corrected to GEOMEAN of reference genes RPLP0 and GUSB. Error bars, s.e.m. Mann-Whitney U test: P < 0.05 (*), P < 0.005 (**), P < 0.0005 (***). (B) Action potential (AP) illustration depicting analyzed electrophysiological parameters. (C) Representative traces of spontaneous APs of day 19 SANCM (blue) and VCM (orange). (D) Cycle length, MDP, APA, Vmax and APD20, APD50 and APD90 of VCM and SANCM at day 19 of differentiation. N =12 cells from 4 independent differentiations. Error bars, s.e.m. Mann-Whitney U test: P < 0.05 (*), P < 0.005 (**), P < 0.0001 (****). (E) Cycle lengths of SANCM and VCM measured at baseline (BL) and after treatment with 3 μM ivabradine (IVA). N = 6 cells from three independent differentiations. Error bars, s.e.m. Wilcoxon: P < 0.05 (*). MDP, maximal diastolic potential; APA, action potential amplitude; Vmax, upstroke velocity; APD20, APD50, APD90, AP duration at 20, 50, 90% repolarization, respectively.

### Unmasking the cellular compositions in SANCM and VCM cultures

The variations in the expression of key genes, such as *TBX18* in SANCM and *MYL2* in VCM are suggestive of heterogeneity in cellular composition (Fig. 1D and 1E). In order to better understand the basis for this, we performed scRNA-seq according to the SORT-seq protocol (Muraro *et al*, 2016). A total of 1287 cells passed pre-processing and quality control. Since plate-to-plate variations were observed, the dataset was corrected using the standard integration workflow on SCTransform normalized data (Fig. S2A) (Hafemeister & Satija, 2019; Stuart *et al*, 2019). Next, unsupervised clustering was performed with the top 15 principal components, which identified 12 clusters. One of the 12 clusters (cluster 9) showed enriched expression of spike-in DNA/ERCCs (Fig S2B), indicating the amplification of mostly ambient RNA and was therefore excluded from further analysis. We also removed two other small clusters (Cluster 10 and 11), which showed enrichment in cell cycle associated genes and genes associated with extraocular muscle development, respectively (Fig. S2B). The remaining 9 clusters (comprised of 1083 cells) were visualized using uniform manifold approximation and projection (UMAP) (McInnes *et al*, 2018) (Fig. 3A). The majority of the clusters highly expressed cardiac sarcomeric genes such as *TNNT2* and *ACTN2*, validating cardiomyocyte identity (Fig. 3B). Clusters containing cells from the VCM differentiation protocol (Clusters 0-3) did not overlap with cell clusters from the SANCM protocol (Clusters 4-8), confirming the generation of transcriptionally different cardiomyocyte subtypes (Fig. S2A).

**Figure 3.**
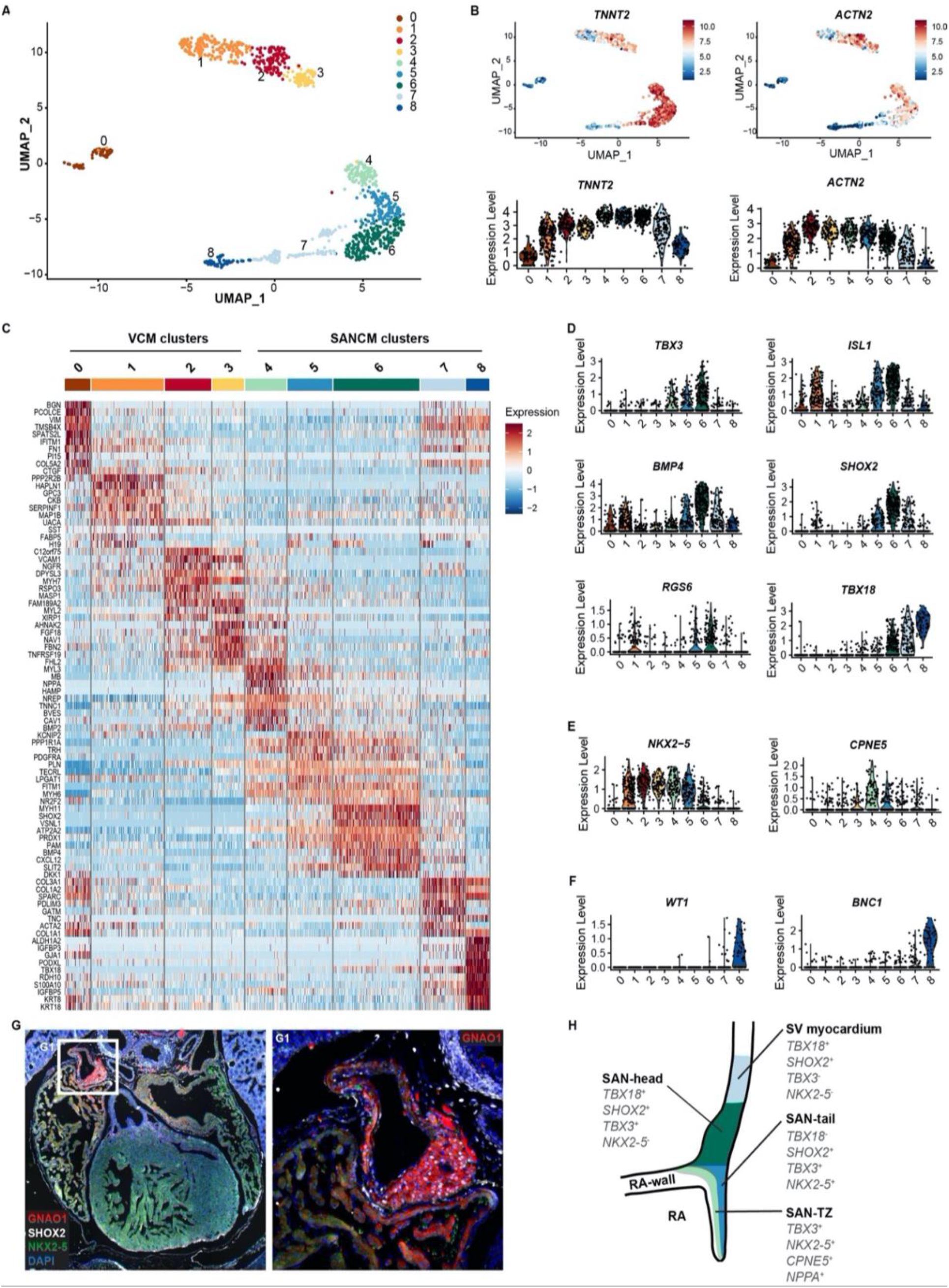
Single cell RNA-sequencing analysis of SANCM and VCM cultures. (A) UMAP representation of cell clusters single cell transcriptomes of SANCM and VCM at day 19 of differentiation. (B) UMAP feature plots and violin plots showing TNNT2 and ACTN2 expression in cell clusters. (C) Heatmap showing the top 10 differentially expressed genes between clusters at day 19 of differentiation. (D-F) Violin plots depicting expression of compact SAN-associated genes (D), SAN-TZ associated genes (E) and proepicaridal associated genes (F). (G) Immunofluorescence staining of GNAO1 co-stained with SHOX2, NKX2-5 and DAPI in E17.5 embryonic mouse heart. G1 is a zoom in of the marked SAN area. (H) Schematic representation of the in vivo organization of the SV and SAN region during development. RA, right atrium; rvv, right venous valve; SAN, sinoatrial node; SV, sinus venosus; UMAP, uniform manifold approximation and projection.

In addition, we observed that the non-cardiomyocyte side populations are specific for each differentiation protocol (Fig. 3A, 3B and S2A). List of genes differentially expressed in each cluster are provided in Table S5.

Analysis of cell clusters belonging to the VCM group unmasked the presence of three cardiomyocyte populations (Cluster 1, 2, 3) and one non-cardiomyocyte population (Cluster 0) as determined by the expression of sarcomeric genes *TNNT2* and *ACTN2* (Fig. 3B). Clusters 2 and 3 expressed *MYH7* and *MYL2* indicating their ventricular identity (Fig. 3C and Fig S2C). However, we observed differences in the expression of other ventricular genes between these two clusters. Whilst the expression of *HOPX* was higher in cluster 2, *HEY2* and *IRX4* expression was restricted to cluster 3 (Fig. S2C). The abundant expression of *HOPX* in cluster 2 likely may represent a more mature cardiomyocyte pool as reported in other similar studies (Churko *et al*, 2018; Friedman *et al*, 2018). Nevertheless, the top 10 differentially expressed genes of cluster 2 and 3 greatly overlap (Fig. 3C) and differences in gene expression may also be the result of different transcriptional states resulting from transcriptional bursts.

Cluster 1 in the VCM group clusters closely with the ventricular cardiomyocytes (Cluster 2 and 3). However, in contrast to the other clusters, it showed lower expression of typical cardiac genes (Fig. 3B). Furthermore, cluster 1 was characterized by the expression of genes such as *HAPLN1*, *GPC3* and *SEMA3C* (Fig. 3C and Fig. S2D), which are associated with progenitors of the myocardial embryonic outflow tract/right ventricle (Sahara *et al*, 2019; Liu *et al*, 2019). An embryonic outflow tract-like cellular identity of cluster 1 is further supported by the expression of *BMP4*, *ISL1* and *PITX2* (Fig. 3D and S2D) and is consistent with previous findings describing co-differentiation of outflow tract-like cells with VCM (Friedman *et al*, 2018). Lastly, the non-cardiomyocyte cluster 0 expressed *NFATC1*, *FOXC1*, *NRG1* and *NPR3* (Fig. S2E), thus representing a fetal endocardial-like lineage (Mikryukov *et al*, 2021).

The SANCM population revealed four cardiomyocyte clusters (Clusters 4–7), marked by *TNNT2* and *ACTN2* expression, and a smaller non-cardiomyocyte cluster (Cluster 8) (Fig. 3A and 3B). Cardiomyocyte clusters 4-6 expressed SAN-associated transcription factors, *TBX3* and *ISL1* as well as *BMP4*, a SAN enriched BMP signaling ligand (van Eif *et al*, 2019), albeit at varying levels (Fig. 3D). However, *SHOX2* and RGS6, encoding a regulator of parasympathetic signaling in heart (Goodyer *et al*, 2019; Yang *et al*, 2010), were restricted to clusters 5 and 6 (Fig. 3D). Moreover, we observed two salient differences between clusters 5 and 6. Whilst cluster 6 expressed *TBX18* besides other key SAN genes, it was devoid of *NKX2-5* (Fig. 3D and 3E), therefore closely resembling the transcriptional signature of the mouse *Tbx18^+^/Nkx2-5^−^* SAN head region (Wiese *et al*, 2009). Cluster 5, on the other hand, revealed a transcriptional pattern found in the SAN tail i.e., *Tbx18^−^/Tbx3^+^/Nkx2-5^+^* (Wiese *et al*, 2009) (Fig. 3D and 3E). The third cardiomyocyte cluster, cluster 4, expressed *TBX3* and lower levels of *ISL1* and *BMP4*, but exhibited higher expression of atrial associated genes, such as *NPPA, HAMP, MYH6* and *NKX2-5* (Fig. 3C, 3E, and S2F), demonstrating that these cells share characteristics of both pacemaker and atrial cells, identified as SAN-TZ cells *in vivo* (Li *et al*, 2019; Goodyer *et al*, 2019). Cluster 4 also revealed higher expression of *CPNE5* (Fig. 3E), which is expressed throughout the entire cardiac conduction system and was found enriched in the transitional SAN region (Goodyer *et al*, 2019). Thus, we determined cluster 4 as SAN-head-like, cluster 5 as SAN-tail-like and cluster 6 as SAN-TZ-like cells.

The remaining two clusters from the SANCM group, clusters 7 and 8, were identified as sinus venosus-like cells and proepicardial cells, respectively. The *TBX18^+^, SHOX2^+^, BMP4^+^, ISL1^+^, TBX3*^-^, *NKX2-5*^−^ expression pattern of cluster 7 (Fig. 3D and 3E) resembles the gene expression pattern of sinus venosus myocardium (Christoffels *et al*, 2006; Blaschke *et al*, 2007; Espinoza-Lewis *et al*, 2009; Cai *et al*, 2003; Vicente-Steijn *et al*, 2010). The expression pattern of cluster 8 was characterized by typical proepicardial markers such as *TBX18, KRT8, KRT18, WT1, BNC1* (Lupu *et al*, 2020) (Fig. 3C, and 3F).

Besides well-established SAN genes, we also identified other markers such as *VSNL1* and *GNAO1*, which were specifically expressed in the SANCM clusters compared with the VCM clusters (Fig. S2H). Visinin like 1 protein (VSNL1, also referred to as VILIP-1 or NVP-1) is a well conserved Ca^2+^ binding protein involved in various cellular signaling cascades (Braunewell & Szanto, 2009) and has previously been identified in mouse, primate as well as human SAN (van Eif *et al*, 2019; Liang *et al*, 2021). Immunofluorescence staining of E17.5 mouse heart confirmed robust expression of VSNL1 in the mouse SAN (Fig. S2I). GNAO1 encodes the guanine nucleotide–binding protein G(o) subunit α, which is a part of the G - protein signal transducing complex (Lambright *et al*, 1994). We corroborated enriched expression of GNAO1 in the SAN of E17.5 mouse (Fig. 3G). Both proteins were also expressed in the atria albeit to a lesser extent (Fig. 3G and S2I).

In summary, hiPSC differentiation towards SANCM closely recapitulates the *in vivo* situation generating subpopulations with gene expression patterns resembling those of SAN-head, SAN-tail and SAN-TZ cardiomyocytes. Furthermore, small populations of co-differentiating sinus venosus-like and proepicardial-like cells alongside SANCM is reflective of shared developmental origins.

### hiPSC differentiation to SANCM recapitulates *in vivo* development

In order to gain a better understanding of the differentiation and specification process of hiPSCs to SANCM, we performed scRNA-seq at several stages during differentiation. At five additional time points (day 0, 4, 5, 6 and 10) (Fig. 4A), cells were sorted and sequenced. A total of 3300 cells including the D19 SANCM population presented in Fig. 3 passed pre-processing and quality control. Unsupervised clustering was performed with the top 20 principal components, which identified 14 clusters. Two of the 14 clusters showed enriched expression of spike-in DNA/ERCCs (Fig. S3A), indicating the amplification of ambient RNA and were therefore excluded from further analysis. The remaining clusters (comprised of 3103 cells) closely correlated with the time of collection, revealing that substantial transcriptional changes occur during the differentiation process *in vitro* (Fig. 4B and S3B). The expression of *TNNT2* and *ACTN2* steadily increased from day 5 (Fig. 4C).

**Figure 4.**
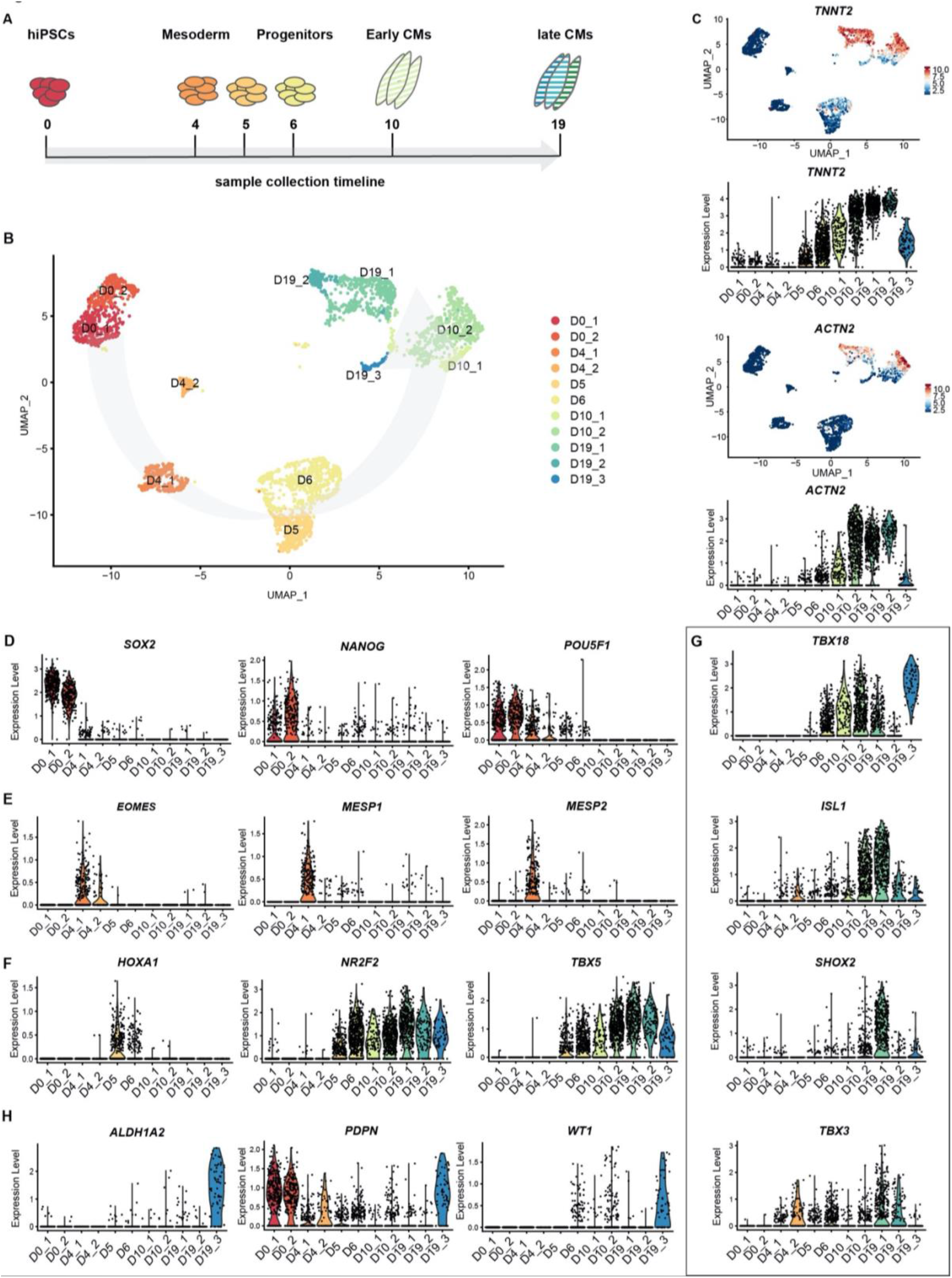
Time course single cell RNA-sequencing of SANCM. (A) Timeline of hiPSC differentiation to SANCM representing sample collection timepoints. (B) UMAP representation of single cell transcriptomes collected at different timepoints throughout differentiation from hiPSC to SANCM. Arrow indicates course of differentiation. (C) UMAP feature plots and violin plots showing TNNT2 and ACTN2 gene expression at different stages of SANCM differentiation. (D-H) Violin plots of pluripotency genes (D), mesodermal genes (E), posterior cardiac progenitor genes (F), SAN-associated transcription factor genes (G) and proepicardial genes (H). hiPSC, human induced pluripotent stem cells; CPC, cardiac progenitor cells; CMs, cardiomyocytes; UMAP, uniform manifold approximation and projection.

Next, we compared the gene expression profile (Table S6) of our time-course dataset with a range of established stage-specific genes reflecting fate choices towards cardiomyocytes. From a pluripotent state at day 0 *(SOX2^+^/NANOG^+^/POU5F1^+^)*, the cells were directed towards germ layer specification with the majority of the cells (Cluster D4_1) exhibiting a cardiac mesoderm-like profile expressing *EOMES*, *MESP1* and *MESP2* (Kitajima *et al*, 2000; Costello *et al*, 2011) (Fig. 4D and 4E). A smaller endoderm-like population was also identified on day 4 (Cluster D4_2), based on the specific expression of *FOXA2* and *SOX17* (Tosic *et al*, 2019) (Fig. S3C). After 24 hours with SAN specification medium (day 5, D5), we identified a gene expression pattern, characteristic for posterior cardiac progenitors (*HOXA1^+^/NR2F2^+^/TBX5^+^)* (Fig. 4F) (Bertrand *et al*, 2011; Stefanovic *et al*, 2020). The first onset of *TBX18* expression was observed at day 6 (D6) of differentiation (Fig. 4G), a transcription factor marking sinus venosus progenitors (Mommersteeg *et al*, 2010).

Early-stage differentiated cells at day 10 (D10) formed two clusters. Cluster D10_1 representing a less mature state compared to cluster D10_2 according to *TNNT2* and *ACTN2* expression (Fig. 4C). Furthermore, cluster D10_1 is partially composed of cells collected on day 19 (D19) of differentiation identified as sinus venosus-like cells (cluster 7) in Fig. 3 (Fig. S3D), suggesting that a fraction of cells was halted during differentiation. Expression of SAN-associated transcription factors *ISL1*, *SHOX2* and *TBX3*, was only obvious at D19 (Fig. 4G). D19 comprised three subpopulations including SANCM (D19_1 and D19_2) and proepicardial-like cells (D19_3), also described in detail in Fig. 3. Interestingly, *PDPN*, reported to be expressed both in the SAN and the epicardium in the mouse heart (Gittenberger-De Groot *et al*, 2007), was found exclusively in the *ALDH1A2^+^/WT1^+^* proepicardial-like population (D19_3) and not in SANCM on day 19 (Fig. 4H).

### WNT signaling mediates the divergence of myocardial and proepicardial lineages

scRNA-seq of SANCM revealed the presence of different SAN subtypes, such as SAN-head, SAN-tail and SAN-TZ, which co-differentiate with a small population of proepicardial-like cells (Epi). To gain insight into the developmental ontogeny of these cell types, we used URD (Farrell *et al*, 2018). URD reconstructs transcriptional trajectories based on user-defined origin (root) and end points (tips). We assigned the cardiac mesoderm stage (day 4) as the root and the distinct subclusters identified on day 19, i.e., SAN-head, SAN-tail, SAN-TZ and proepicardial cells as the tips, resulting in a pseudotime tree consisting of six main segments (Fig. 5A). Sinus venosus-like cells were excluded as a tip since it partially clustered with progenitors of day 10 and is a cell type independent of the SAN niche (Fig. S3D). Cells from day 5, day 6 and a fraction of day 10 were located near the root of the tree in segment 1, constituting a common progenitor pool. From segment 1, the pseudotime tree branches off into two lineages, the proepicardial branch (Segment 2) and the myocardial branch (Segment 3). The proepicardial branch contained cells collected on day 10 as well as day 19, whereas the myocardial branch primarily consisted of cells collected on day 10. Whilst most myocardial cells at day 10 were present in segment 3, a small fraction appeared committed to SAN-TZ lineage (Segment 4). SAN-tail (Segment 5) and SAN-head (Segment 6) were assigned later pseudotimes and only contained cells from day 19.

**Figure 5.**
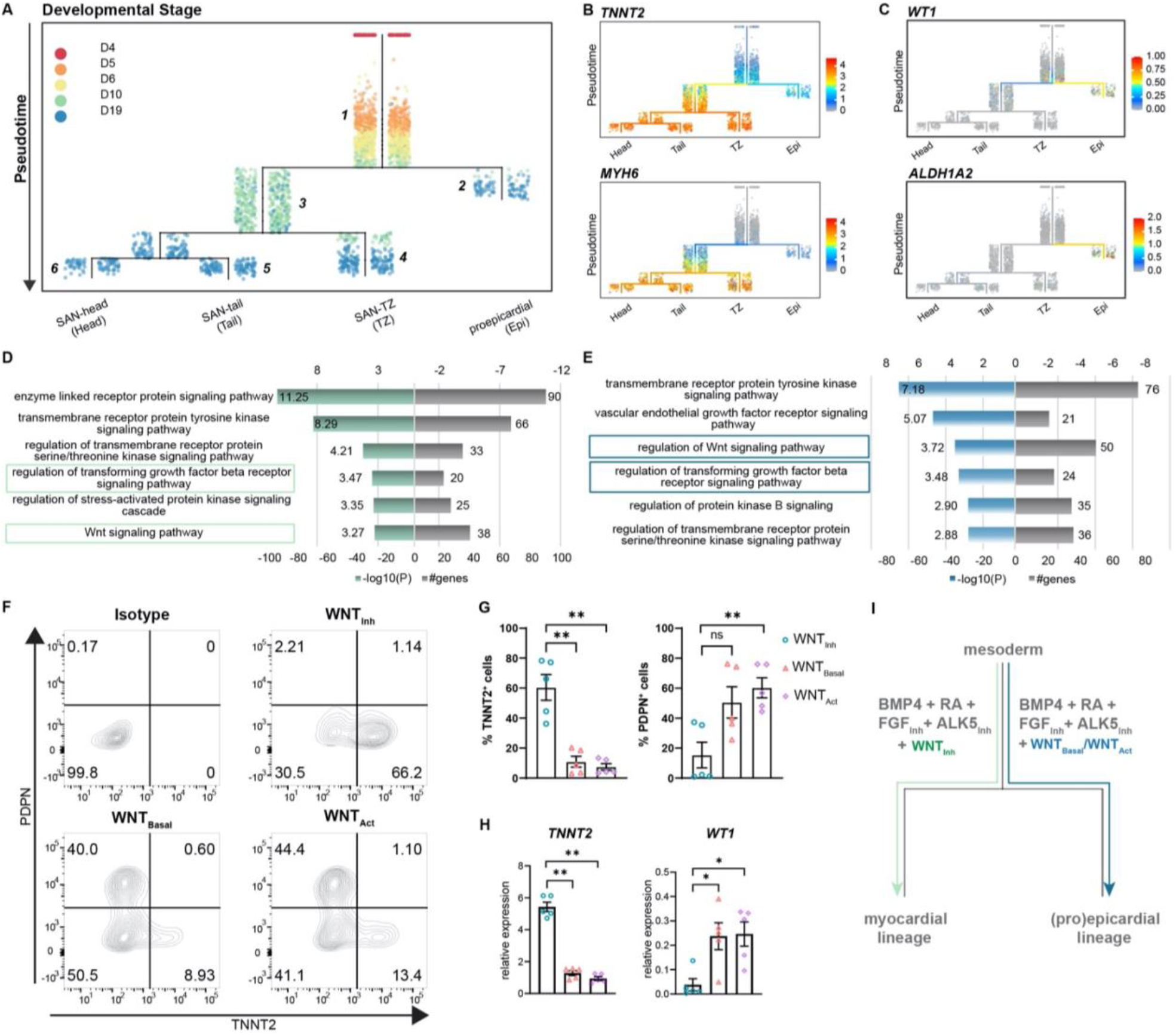
Reconstruction of single cell trajectories. (A) URD trajectory tree starting at the mesodermal stage (day 4) and proceeds to the terminally differentiated cell clusters identified on day 19. Colors correspond to the timepoint of cell collection. (B) Expression of TNNT2 and MYH6 marking the myocardial lineage (C) Expression of WT1 and ALDH1A2 marking the proepicardial lineage in the trajectory tree. (D-E) Representative GO terms based on differentially expressed genes between the common progenitor, segment 1 and the myocardial branch, segment 3 (D) or the proepicardial branch, segment 2 (E) lineage. (F) Representative contour plots and (G) summarized data demonstrating percentage of TNNT2^+^ and PDPN^+^ cells in baseline condition containing WNT inhibitor, XAV (WNT_Inh_), excluding WNT inhibitor, XAV (WNT_Basal_) and addition of WNT activator, CHIR (WNT_Act_). N = 5 independent differentiations. Error bars represent s.e.m., Kruskal-Wallis, post-hoc Mann-Whitney U test: P < 0.005 (**). (H) RT-qPCR demonstrating the expression of caridomyocyte gene TNNT2 and the proepicaridal gene WT1 in WNT_Inh_, WNT_Basal_ and WNT_Act_ conditions. N= 5 independent differentiations; corrected to GEOMEAN of reference genes RPLP0 and GUSB. Error bars, s.e.m. Kruskal-Wallis, post-hoc Mann-Whitney U test: P < 0.05 (*), P < 0.005 (**). (I) Schematic representation of divergence of SAN myocardial and proepicardial lineages from a common progenitor.

The ordering of cell populations in the trajectory tree suggests that cells on day 5 and day 6 could potentially give rise to both the myocardial and proepicardial lineages. The first divergence was only apparent at day 10 (Fig. 5A) with the majority of the cells directed towards the myocardial lineage whereas a small population branched off towards the proepicardial lineage. Accordingly, cardiomyocyte genes such as *TNNT2* and *MYH6* were selectively expressed in the myocardial branch (Fig. 5B), and proepicardial genes such as *WT1* and *ALDH1A2* were enriched in the proepicardial branch (Fig. 5C). Thus, day 10 of differentiation appears to be a critical branching point for myocardial and proepicardial cell fates driven by BMP and RA.

In order to identify the key players that regulate myocardial versus proepicardial cell fate, we performed gene ontology (GO) analysis on differentially expressed genes between the common progenitor (Segment 1) and the myocardial branch (Segment 3) or the common progenitor (Segment 1) and the proepicardial branch (Segment 2) (Fig. 5A). GO term analysis identified signaling pathways potentially involved in myocardial versus proepicardial divergence (Table S7). As both groups contained a number of genes implicated in TGFβ and WNT signaling (Fig. 5D and 5E), we tested the impact of manipulating these signaling pathways on myocardial versus proepicardial fate specification. The standard SANCM differentiation cocktail contains the ALK5 inhibitor (SB431542) included to offset any effects of BMP4 on TGFβ signaling (Birket *et al*, 2015; Protze *et al*, 2017). To allow active TGFβ signaling, we excluded the ALK5 inhibitor (w/o ALK5_inh_) from the SANCM differentiation cocktail. Our results show that active TGFβ signaling does not alter myocardial vs proepicardial cell fate, as determined by the expression of TNNT2 marking cardiomyocytes, and PDPN marking proepicardial cells, in the resulting population (Fig. S4A and S4B).

Next, we tested the role of WNT signaling in myocardial versus proepicardial branching. From the SANCM differentiation cocktail containing the WNT inhibitor, XAV939 (WNT_Inh_) (Fig. 1A), we either removed the WNT inhibitor (WNT_Basal_), or replaced it with the WNT agonist, CHIR (WNT_Act_). Applying the standard cocktail containing the WNT inhibitor resulted in an average of 60% TNNT2^+^ cells with a small side population of 10% PDPN^+^ cells (Fig. 5F and 5G). Strikingly, removal of the WNT inhibitor strongly compromised the percentage of cardiomyocytes (~10% TNNT2^+^), whereas percentage of PDPN^+^ cells increased. Addition of a WNT agonist had a similar effect although it did not further enhance the percentage of proepicardial cells. RT-qPCR confirmed a proepicardial-like gene expression in WNT_Basal_ and WNT_Act_ conditions evidenced by higher expression of *WT1* and lower expression *TNNT2* mRNA compared with WNT_Inh_ (Fig. 5H). These findings demonstrate that in the presence of active WNT signaling, BMP and RA steer common progenitors towards the proepicardial fate and that inhibition of WNT signaling is crucial for their differentiation towards the myocardial lineage (Fig. 5I).

### Diversification between the myocardial SAN subpopulations involves TGFβ and WNT signaling

Members of the TGFβ/BMP signaling pathway were preferentially expressed in SANCM subpopulations at day 19. These included ligands *BMP4* (SAN-head) and *BMP2* (SAN-TZ), as well as genes involved in negative regulation of TGFβ/BMP signaling such as *HTRA1* (SAN-head; SAN-tail) and *FBN2* (SAN-TZ) (Fig. 6A), implicating TGFβ/BMP signaling in differentiation towards SAN subpopulations. Moreover, the percentage of TNNT2^+^ cells in the w/o ALK5_inh_ condition were unaffected (Fig. S4A and S4B) suggesting that active TGF β signaling does not affect myocardial specification itself. These observations led us to evaluate the identity of cardiomyocytes obtained in w/o ALK5_Inh_ condition. To identify the effect of removing the ALK5 inhibitor from the SANCM differentiation cocktail on SAN subpopulations, we assessed the expression of genes specific to or enriched in individual SAN fractions, such as *SHOX2*, *VSNL1*, *NTM* and *FLRT3* for SAN-head, *KCNIP2* for SAN-tail and *NKX2-5*, *NPPA* and *CPNE5* for SAN-TZ (Fig. S5A). Gene expression analysis of the resulting population at day 19 of differentiation revealed increased expression of tail and TZ-enriched genes, such as *KCNIP2* and *NPPA* (Fig. 6B). Other SAN-TZ enriched genes, such as *CPNE5* and *NKX2-5*, also showed a trend for higher expression, but were not statistically different (Fig. 6B). Furthermore, we did not observe significant differences in the expression of genes associated with SAN-head (Fig. S5B).

**Figure 6.**
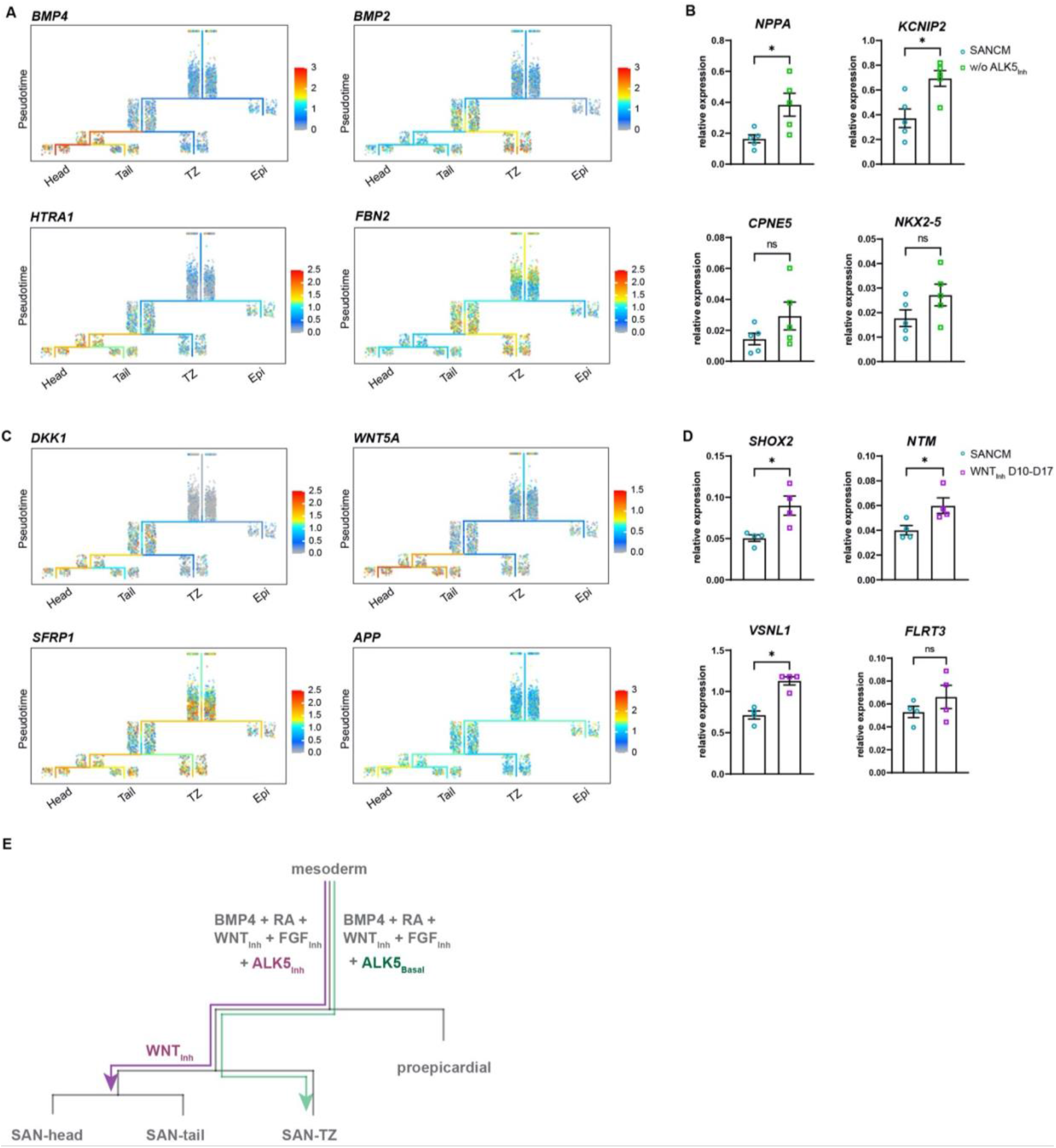
Diversification of SAN-head, SAN-tail and SAN-TZ subpopulations. (A) Expression of TGFβ signaling pathway members in the trajectory tree. (B) RT-qPCR of genes enriched in the SAN-TZ lineage upon exclusion of ALK5 inhibitor from baseline condition (w/o ALK5Inh). N= 5 independent differentiations; corrected to GEOMEAN of reference genes RPLP0 and GUSB. Error bars, s.e.m. Mann-Whitney U test: P < 0.05 (*). (C) Expression of WNT signaling pathway members in the trajectory tree. (D) RT-qPCR of genes enriched in the SAN-head lineage upon prolonged WNT signaling inhibition (XAV D10-17). N= 4 independent differentiations; corrected to GEOMEAN of reference genes RPLP0 and GUSB. Error bars, s.e.m. Mann-Whitney U test: P < 0.05 (*). (E) Schematic representation of the diversification of the various SANCM subpopulations from a common myocardial progenitor.

In order to better understand the mechanisms implicated in SAN-head differentiation, we looked at transcriptional changes between common progenitor state (Segment 3) and day 19 SANCM subpopulations (Segment 4-6). Ordering of cells in the trajectory tree in Fig. 5A suggests that a large majority of myocardial cells remain uncommitted at day 10 and specification towards SAN-head and SAN-tail cells only occurs after day 10. GO term analysis of differentially expressed genes between the common myocardial progenitor at day 10 (Segment 3) and each SANCM subtype of day 19 (Segment 4-6) revealed enrichment of several WNT signaling modulators, such as *DKK1*, *WNT5A*, *SFRP1* and *APP*, primarily in the SAN-head branch (Segment 6) (Fig. 6C and Table S8). Whilst DKK1 is an inhibitor of canonical WNT signaling, WNT5A is a non-canonical WNT ligand. Therefore, we posited that inhibition of canonical WNT signaling may enhance differentiation to SAN-head. To test this assumption, we treated SANCM cultures with XAV 939 from day 10 to day 17, following the findings from the trajectory tree. Inhibition of canonical WNT signaling from day 10–17 significantly increased expression of SAN-head enriched genes, such as *SHOX2*, *VSNL1* and *NTM*, and a trend for higher expression in *FLRT3*, but did not influence the expression of SAN-tail or SAN-TZ genes (Fig. 6D and S5C). Taken together, these findings underscore a stage-specific role for TGFβ and WNT signaling in differentiation towards specific SANCM subpopulations.

## Discussion

The developmental ontogeny of the SAN is poorly understood. Here, we aimed to study the differentiation and diversification of the human SAN *in vitro*. Directed differentiation of hiPSCs to cardiomyocytes was achieved using a two-step approach, wherein hiPSCs were first directed towards mesoderm, which was further steered towards a cardiac fate by inhibition of WNT signaling. This standard approach generated cardiomyocytes with a ventricular-like signature. Alongside inhibition of WNT signaling, addition of BMP4 and RA as well as ALK5 inhibitor, and FGF inhibitor, at the cardiac mesoderm stage resulted in cardiomyocytes with SAN-like profile as reported before (Protze *et al*, 2017). Whereas the identity of SAN cells obtained in this previous study appears to be predominantly SAN-head-like inferred by NKX2.5^−^ and TNNT2^+^ expression, we found the presence of multiple cell types that develop at the inflow tract of the heart in our *in vitro* cultures. Whether this is due to differences in the methods used for mesoderm induction or due to different culture conditions (3D vs 2D) is not known.

Current differentiation protocols for SANCM result in 25 – 50% NKX2-5^−^ SANCM resembling SAN-head (or even a fraction of sinus venosus-like cells) while no data is available w.r.t the presence of NKX2.5^+^ SAN-tail and SAN-TZ cells (Protze *et al*, 2017; Ren *et al*, 2019). We found that 54% of the entire SANCM pool exhibited the expression pattern described in previous studies (*NKX2-5^−^/TNNT2^+^*). A fraction of these NKX2-5^−^ cardiomyocytes (18% of total SANCM pool) resembled the gene expression pattern of sinus venosus myocardium, characterized by lower expression of sarcomeric genes such as *TNNT2* and *ACTN2*, presence of *SHOX2* and absence of *TBX3*. Importantly, 36% of the total SANCM pool revealed a gene expression pattern similar to the SAN-head (*NKX2-5^−^/TBX18^+^/TBX3^+^*), whereas 19% resembled a SAN-tail like phenotype (*NKX2-5^+^/TBX18^−^/TBX3^+^*) and 17%, a transitional cell like phenotype (*NKX2-5^+^/CPNE5^+^/TBX3^+^*) (Wiese *et al*, 2009; Sizarov *et al*, 2011; Goodyer *et al*, 2019).

To gain insight into the origin and diversification of the cell types in the SANCM group, we applied trajectory inference analysis. Our data revealed early divergence between the myocardial SAN and proepicardial populations in line with previous reports that identified a *Tbx18^+^* common progenitor for these lineages (van Wijk *et al*, 2009; Kruithof *et al*, 2006). Besides SAN development, BMP and RA signaling are also implicated in the development of the epicardium and a cross talk with WNT signaling has been postulated (Wiesinger *et al*, 2021). Our results show that WNT signaling in fact determines the bifurcation of myocardial and proepicardial cell fates. Excluding the WNT inhibitor in the presence of BMP4 and RA, diminished the myocardial population and enriched the proepicardial population. Our findings also correlate with a previous study, which described the generation of proepicardial cells under similar experimental conditions (Guadix *et al*, 2017). Comparably, the PDPN^+^ proepicardial population in the study of Guadix et al was prominent in culture condition with BMP4 and RA, which strongly decreased with the addition of a WNT inhibitor. Active WNT signaling therefore seems pertinent for epicardial cell differentiation, as indicated in several *in vitro* differentiation studies (Witty *et al*, 2014; Iyer *et al*, 2015; Bao *et al*, 2016; Zhao *et al*, 2017). Consistent with our results, intrinsic WNT activity is sufficient for epicardial differentiation by BMP4 and RA (Iyer *et al*, 2015; Guadix *et al*, 2017). Similarly, activation of WNT signaling at the *NKX2.5*^+^ cardiac progenitor stage resulted in SAN cardiomyocytes as well as a non-cardiomyocyte population, which exhibited an epicardial like phenotype (Ren *et al*, 2019).

Our time series dataset further provided insight into the developmental trajectory of SANCM. A proportion of the cardiac progenitor cells collected on day 10 were already committed to the SAN-TZ lineage, whereas none of these progenitors appeared determined to SAN-head or SAN-tail lineages indicating that SAN-TZ cell specification occurs earlier. The order of differentiation of the components of the mouse sinus venosus and SAN have been analyzed in detail. During mouse caudal heart development, *Tbx18^−^* posterior second heart field progenitors first form the inflow tract of the myocardial heart tube, which differentiate into atrial cardiomyocytes. Subsequently, *Tbx18^+^* progenitors differentiate to cardiomyocytes and form the SAN and sinus venosus components in the order of their future anatomical position from proximal to distal of the atrial myocardium. Thus, the *Tbx18^+^* progenitors first form the *Tbx3* transitional pacemaker cells, directly followed by the *Tbx3^+^* SAN tail, the *Tbx3^+^* SAN head and finally the *Tbx3^−^* sinus venosus myocardium of the superior caval vein (Christoffels *et al*, 2006; Mommersteeg *et al*, 2007; Wiese *et al*, 2009; Mommersteeg *et al*, 2010; Mohan *et al*, 2018). It is therefore interesting to note that the developmental trajectory of the SANCM cells *in vitro* recapitulate this temporal aspect of *in vivo* mouse SAN development. A fraction of the cells collected on day 19 formed a small common segment before separating into the SAN-head or the SAN-tail tips indicating that differentiation of the cell types may not yet be complete. Nevertheless, our data shows that SAN-head, SAN-tail and SAN-TZ originate from a common progenitor, which under the influence of various signaling pathways diversify into these subpopulations.

Consistent with our findings, TGFβ/BMP signaling mediators have been found enriched in the embryonic SAN (Vedantham *et al*, 2015; van Eif *et al*, 2019; Li *et al*, 2019; Goodyer *et al*, 2019), which is maintained in adulthood (Linscheid *et al*, 2019; Brennan *et al*, 2020). Even though the exact role of TGFβ/BMP signaling during SAN development is not known, it has been proposed to be involved in recruitment of proepicardial cells and remodeling of interstitium in the SAN niche (Easterling *et al*, 2021). Comparably, WNT signaling has been described as a critical cue for SAN development (Bressan *et al*, 2013; Ren *et al*, 2019). Our results reveal a role for TGFβ and WNT signaling in enhancing gene signature pertaining to SAN-TZ and SAN-head, respectively. These changes do not seem to occur at the expense of other subpopulations and further studies are needed to evaluate the role of additional signaling pathways and combinatorial modulations that precisely alter cell fate of the common progenitor.

Principles of stage-specific manipulation of signaling pathways described in this study can be applied to other pluripotent stem cell lines including patient-specific lines to obtain desired cell fractions. Derivation of SAN-head, SAN-tail and SAN-TZ cells *in vitro* will allow us to perform a comparative assessment of the functional properties of these cell types. Relatively little is known regarding the role of SAN-tail and SAN-TZ cells in impulse initiation and propagation. Furthermore, engineered tissues comprised of these cell types will enable modeling of complex diseases such as SAN exit block, which occurs due to impaired impulse propagation to the atria and is thought to result from dysfunctional transitional cells(Li *et al*, 2019). Importantly, incorporation of SAN subpopulations in the design of biomimetic cell constructs would permit the evaluation of optimal configurations that effectively regenerate the dysfunctional pacemaker tissue.

## Materials and Methods

### Maintenance of hiPSC lines and differentiation to CMs

hiPSC line LUMC0099iCTRL04 used in this study was generated by the iPSC core facility of Leiden University Medical Center following due protocols for informed consent and use of these cells for research purposes. The cell line is registered in Human Pluripotent Stem Cell Registry (https://hpscreg.eu/cell-line/LUMCi004-A).

hiPSCs were maintained in mTESR1 medium (Stem cell technologies, #5850) on growth factor reduced Matrigel (Corning, #356234) at 37° C with 5% CO_2_ and passaged once a week. For cardiac differentiation, cells were seeded at a density of 2.5-3×10^4^ cells/cm^2^. Differentiation was induced when cells reached 80-90% confluency using BPEL medium (Ng *et al*, 2008) supplemented with 20 ng/mL Activin-A (Miltenyi Biotec, #130-115-012), 20 ng/mL BMP4 (R&D systems, #314-BP-010/CF), and 1.5 μmol/L CHIR99021 (Axon Medchem, #1386). Three days after initiation, medium was replaced with BPEL containing 5 μmol/L XAV939 (Tocris Bioscience, #3748/10). For SANCM differentiation, 5 μmol/L XAV939, 2.5 ng/mL BMP4, 5 μmol/L SB431542 (Tocris, #1614), 250 nmol/L RA (Sigma, #R2625-50MG) and 250 nmol/L PD173074 (Selleck Chemicals, #1264) were added on day 4. Differentiation medium was replaced with BPEL medium after 48 hours (SANCM) or 96 hours (VCM) and cells refreshed every three days thereafter. To evaluate the role of canonical WNT signaling for differentiation towards SAN-head lineage, XAV939 (5 μmol/L), was added from day 10 – day 17.

### RT-qPCR

Total RNA of day 19 hiPSC-derived cultures was isolated using Nucleospin RNA kit (Machery Nagel, # MN740955.50) according to the manufacturer’s instructions. Reverse transcription was performed using Superscript II (ThermoFisher Scientific, #18064071) with oligo dT primers (125 μmol/L). qPCR was performed on the LightCycler 2.0 Real-Time PCR system (Roche Life Science). Primer pairs were designed to span an exon-exon junction or at least one intron (Table S3). qPCR reaction mix was prepared using the LightCycler 480 SYBR Green I Master (Roche, #4887352001), primers (1 μmol/L) and cDNA (equivalent to 10 ng RNA). Amplification of target sequences was performed using the following protocol: 5 min at 95 °C followed by 45 cycles of 10 sec at 95 °C, 20 sec at 60 °C, and 20 sec at 72 °C. Data analysis was performed using LinRegPCR program (Ruijter *et al*, 2009). For data normalization, two experimentally assessed reference genes, *RPLP0* and *GUSB* were used.

### Immunofluorescence staining

Cells cultured as a confluent monolayer on glass coverslips were fixed with 4% paraformaldehyde. Permeabilization was performed with 0.1% Triton-X (Sigma Aldrich #T8787) and a blocking step was carried out with 4% swine serum (Jackson Immunoresearch, #014-000-121) for one hour. Primary and secondary antibodies were diluted in 4% swine serum as stated in Table S4 and incubated at room temperature for 1 hour or at 4 °C overnight. Cell nuclei were stained with DAPI (Sigma Aldrich #D9542). Imaging was carried out with Leica TCS SP8 X DLS confocal microscope. Data visualization and processing was performed with the Leica LAS-X software.

### Single Cell Patch-Clamp

Cardiomyocytes were dissociated using 1x TryPLE Select (ThermoFisher Scientific #12563011) and plated at a density of 7.0^10^3^ per coverslip. After one week, cells with a smooth surface and intact membrane were chosen for measurements. Action potentials were recorded at 37°C with the amphotericin-B-perforated patch-clamp technique using a Axopatch 200B Clamp amplifier (Molecular Devices Corporation). Measurements were carried out in Tyrode’s solution containing 140 mmol/L NaCl, 5.4 mmol/L KCl, 1.8 mmol/L CaCl2, 1.0 mmol/L MgCl2, 5.5 mmol/L glucose and 5.0 mmol/L HEPES. pH was adjusted to 7.4 with NaOH. Pipettes (borosilicate glass; resistance 1.5-2.5 MΩ) were filled with a solution containing 125 mmol/L potassium gluconate, 20 mmol/L KCl, 10 mmol/L NaCl, 0.4 mmol/L amphotericin-B and 10 mM HEPES, pH was adjusted to 7.2 with KOH. Signals were low-pass-filtered (cut off frequency 10 kHz) and digitized at 40 kHz. Action potentials were corrected for the estimated change in liquid junction potential (Barry & Lynch, 1991). Data acquisition and analysis were performed using custom software.

### Immunohistochemistry on mouse heart tissue

Paraffin-embedded hearts were sectioned at 7 uM. Sections were mounted onto silane-coated slides, deparaffinized in xylene, rehydrated in graded ethanol series and washed in phosphate-buffered saline (PBS, pH 7.4). Heat-induced antigen retrieval was performed using unmasking solution (Vector Labs #H-3300-250). Sections were incubated with primary antibodies (Table S4) diluted in 4% bovine serum albumin (BSA; Sigma Aldrich #A7906) at 4 °C overnight. After washing in TBST buffer (25 mM Tris, 150 mM Nacl, 2.5 mM Kcl and 0.5% Tween w/v) sections were incubated with fluorochrome-conjugated secondary antibodies at room temperature for 2 hours in the dark. Sections were washed in TBST, stained with DAPI (Sigma Aldrich #D9542) and mounted in PBS-Glycerol (1:1). Imaging was performed with Leica DMI6000 inverted microscope.

### Flow cytometry

Cardiomyocytes were dissociated using 1x TrypLE Select (ThermoFisher Scientific #12563011). For intracellular staining, cells were fixed and stained using the FIX & PERM kit (Thermofisher; # GAS004) according to the manufacturer’s instructions. For cell surface antigens, the antibody (Table S4) was added to the cell suspension resuspended in a buffer containing 10% BSA (Sigma Aldrich, #A8022) and 0.5M EDTA (ThermoFisher Scientific #15575020). All antibody incubations were performed for 30 minutes on ice protected from light. Acquisition was performed on FacsCanto II Cell Analyzer (Beckton Dickinson). Data was analyzed using FlowJo v10. Antibody information is provided in Supplementary Table 6.

### Cell sorting for single cell RNA sequencing

Single cell sequencing was performed using SORT-seq method (Muraro *et al*, 2016). Cells from one representative differentiation were collected at different stages (day 0, 4, 5, 6 and 10). At the end time point on day 19, cells from two independent differentiations were collected to ascertain reproducibility. For each time point, cells were sorted into two 384-well plates, each well containing an oil droplet with barcoded primers, spike-ins and dNTPs. Preparation of single-cell libraries was performed using the CEL-Seq2 protocol (Muraro *et al*, 2016; Hashimshony *et al*, 2016). Paired-end sequencing was performed on the NextSeq500 platform using 1×75 bp read length kit.

### Bioinformatic Analysis

#### Reference genome annotation

Mapping was performed using BWA-MEM against the (human) genome assembly GRCh38 (hg38). Count matrices were generated using MapAndGo, filtering reads with a minimum quality score of 60 and no alternative hits.

#### scRNAseq data preprocessing, normalization and batch-correction, clustering, differential gene expression, cell-type identification and visualization

Data analysis was performed using the R toolkit Seurat version 3 (Stuart *et al*, 2019). Data QC and preprocessing, dimensional reduction, clustering and differential gene expression were performed according to the standard workflow (https://satijalab.org/seurat/). Briefly, high quality single cells collected on D19 were selected according to the following parameters: gene count > 1,000 and < 9,000, mRNA molecule count < 60,000 and mitochondrial gene count < 50%. The filters for the time series dataset were set the following: gene count > 600; mRNA molecule count < 100,000; mitochondrial gene count < 50%. Next, normalization, scaling and identification of variable features (nfeatures=3000) based on variance stabilizing transformation (“vst”) was performed using the SCTransform command (Hafemeister & Satija, 2019). Since technical plate-to-plate variations were observed, SCTransform data integration was performed by normalizing each dataset individually, identifying integration anchors within the datasets collected on the same timepoint and integrating the datasets. Dimensionality reduction was performed using principle component (PC) analysis and Uniform Manifold Approximation and Projection (UMAP) with the top 15 PCs (day 19 dataset) or top 20 PCs (day 0 - 19 SANCM dataset) and seed set to 2020. For cell clustering, a KNN (K-nearest neighbor) graph was constructed based on euclidean distance in PCA space and clusters were identified using the Louvain algorithm, as implemented in the FindNeighbors and FindClusters command. Identified clusters were then visualized in a UMAP using the DimPlot command. For differential expression testing and visualization, LogNormalization was performed according to the standard workflow on the uncorrected dataset and differential gene expression was determined using Wilcoxon rank sum test. Differentially expressed gene lists show genes, which are expressed in at least 25% in either of the two fractions of cells and limited to genes, which are differentially expressed (on average) by at least 0.25-fold (log-scale) between the two compared cell fractions. Cell type specific marker genes were used to annotate cell clusters. VlnPlot, FeaturePlot and DoHeatMap commands were used to visualize gene expression.

#### Pseudotime and trajectory inference

For the reconstruction of transcriptional trajectories from the mesodermal stage (day 4) to SANCM (day 19), the URD algorithm was used (Farrell *et al*, 2018). hiPSC clusters (D0_1, D0_2) were excluded as we reasoned that cell lineage diversification will not occur before mesoderm induction. A small endoderm-like cluster (D4_2) was also excluded. Identification of highly variable genes, PCA and tSNE projection (RunTSNE command, dims=1:20) were performed using Seurat, as described above. The Seurat object was converted into a URD object. All steps were performed according to the manual provided by the Schier lab (https://schierlab.biozentrum.unibas.ch/urd).

Briefly, K-nearest neighbor graph was calculated using k=100 and poorly connected cells (outliers) were removed. Outliers were identified as cells, which are unusually far from their nearest neighbor and their 20^th^ nearest neighbor (based on their distance to their nearest neighbor). Next, transition probabilities were calculated between transcriptomes to connect cells with similar gene expression patterns and a diffusion map was constructed using knn=50 and global sigma=12. Diffusion map was visualized and assessed by plotting diffusion component pairs using PlotDimArray function. Then, the root of the specification tree was defined (cells in cluster D4_1, corresponding to mesoderm stage) and pseudotime was assigned to each cell by simulated “floods” (n=100, minimum.cells.flooded = 2), using previously calculated transition probabilities. The tips of the trajectory tree were assigned using clusters derived from terminally differentiated cells (day 19). Cluster 7 (Fig. 3A) was not assigned as tip cluster as those cells appear to be halted during differentiation. Cluster 4, 5, 6 and 8 were used as tip clusters corresponding to SAN-TZ, SAN-tail, SAN-head and proepicardial like cells, respectively. Trajectories from the tips back to the root were identified using biased random walks with the following parameters: optimal.cells.forward=50, max.cells.back=80; n.per.tip = 25000, root.visits = 1, max.steps = 5000. In order to build the developmental trajectory and branching tree structure, the visitation frequency of each cell was determined by the random walks from each tip. Visitation frequencies were visualized to ensure a well-connected tree structure from the tips to the root. Lastly, the branching tree structure was constructed using the following parameters: divergence.method = “preference”, cells.per.pseudotime.bin = 35, bins.per.pseudotime.window = 10, save.all.breakpoint.info = T, p.thresh=0.000001. Gene expression within the dendrogram was visualized using the plotTree command. Differential gene expression between different segments of the developmental tree were performed using the markersAUCPR command (auc.factor = 0.9, effect.size = 0.4, frac.must.express = 0.5).

### Gene ontology enrichment analysis

Gene ontology enrichment analysis was performed using Protein Analysis Through Evolutionary Relationships (PANTHER) Classification System version 16.0, release date 2020-12-01 (Ashburner *et al*, 2000; Carbon *et al*, 2021).

### Statistical analysis

Statistical analysis was carried out in GraphPad Prism version 9.1.0 for Windows GraphPad Software, San Diego, California USA, www.graphpad.com. Data were represented as mean ± standard error of the mean (s.e.m.). Non-parametric tests were performed in all cases. Number of samples (n) and the method used to test statistical significance are stated in each figure legend. *P* < 0.05 was considered statistically significant.

## Data and Code Availability

Original sequencing data has been deposited in NCBI GEO repository. Data will be publicly available after peer-review and formal publication of the study. R scripts will be available upon request.

## Acknowledgements

We thank Berend Hooibrink from the Flow Cytometry facility, Department of Medical Biology, for assistance with cell sorting and Corrie de Gier-de Vries, Department of Medical Biology, for help with histology of mouse hearts.

We also gratefully acknowledge funding from the European Research council starting grant 714866 and associated proof-of-concept grant 899422, Health Holland LentiPace II, Horizon 2020 Eurostars (E114245 and E115484), Dutch Research Council Open Technology Program 18485 to GJJB; Netherlands Organization for Health Research and Development (ZonMW), ZonMW TOP 40-00812-98-17061 to VMC, ZonMW and the Dutch Heart foundation MKMD grant 114021512 and Dutch Heart Foundation Dekker fellowship 2020T023 to HDD.

## Author Contributions

Conceptualization, data collection, interpretation and writing A.W and H.D.D; scRNA-seq data analysis and visualization A.W; Data collection and analysis J.L, L.F, P.B, A.O.V; Critical feedback, discussion, and resources, V.M.C, G.J.J.B; Review and Editing, V.M.C, G.J.J.B, A.O.V; Funding acquisition, G.J.J.B and H.D.D. Supervision, H.D.D. All authors read and approved the final manuscript.

## Supplementary Material

### Data Supplement

Supplementary tables S1–S4 and Supplementary figures S1–S5.

### Supplementary Excel File

**Table S5.** Differentially expressed genes in SANCM and VCM clusters identified at day 19. Related to Figure 3 and S2.

### Supplementary Excel File

**Table S6.** Differentially expressed genes in clusters identified in time series analysis. Related to Figure 4 and S3.

### Supplementary Excel File

**Table S7.** GO term analysis of differentially expressed genes between common progenitor segment (1) and proepicardial segment (2) or myocardial segment (3). Related to Figure 5.

### Supplementary Excel File

**Table S8.** GO term analysis of differentially expressed genes between myocardial (3) and SAN-TZ segment (4) or SAN-tail segment (5) or SAN-head segment (6). Related to Figure 6.

## References

Ashburner M, Ball CA, Blake JA, Botstein D, Butler H, Cherry JM, Davis AP, Dolinski K, Dwight SS, Eppig JT, et al (2000) Gene Ontology: tool for the unification of biology. Nature Genetics 25: 25–29

Bao X, Lian X, Hacker TA, Schmuck EG, Qian T, Bhute VJ, Han T, Shi M, Drowley L, Plowright AT, et al (2016) Long-term self-renewing human epicardial cells generated from pluripotent stem cells under defined xeno-free conditions. Nature Biomedical Engineering 1

Barry PH & Lynch JW (1991) Liquid junction potentials and small cell effects in patch-clamp analysis. The Journal of Membrane Biology 121: 101–117 doi:10.1007/BF01870526 [PREPRINT]

Bertrand N, Roux M, Ryckebüsch L, Niederreither K, Dollé P, Moon A, Capecchi M & Zaffran S (2011) Hox genes define distinct progenitor sub-domains within the second heart field. Developmental Biology 353: 266–274

Birket MJ, Ribeiro MC, Verkerk AO, Ward D, Leitoguinho AR, Den Hartogh SC, Orlova V V., Devalla HD, Schwach V, Bellin M, et al (2015) Expansion and patterning of cardiovascular progenitors derived from human pluripotent stem cells. Nature Biotechnology 33: 970–979

Blaschke RJ, Hahurij ND, Kuijper S, Just S, Wisse LJ, Deissler K, Maxelon T, Anastassiadis K, Spitzer J, Hardt SE, et al (2007) Targeted mutation reveals essential functions of the homeodomain transcription factor Shox2 in sinoatrial and pacemaking development. Circulation 115: 1830–1838

Boyett MR, Honjo H & Kodama I (2000) The sinoatrial node, a heterogeneous pacemaker structure. Cardiovascular Research 47: 658–687 doi:10.1016/S0008-6363(00)00135-8 [PREPRINT]

Braunewell KH & Szanto AJK (2009) Visinin-like proteins (VSNLs): Interaction partners and emerging functions in signal transduction of a subfamily of neuronal Ca2+-sensor proteins. Cell and Tissue Research 335: 301–316 doi:10.1007/s00441-008-0716-3 [PREPRINT]

Brennan JA, Chen Q, Gams A, Dyavanapalli J, Mendelowitz D, Peng W & Efimov IR (2020) Evidence of Superior and Inferior Sinoatrial Nodes in the Mammalian Heart. JACC: Clinical Electrophysiology 6: 1827–1840

Bressan M, Henley T, Louie JD, Liu G, Christodoulou D, Bai X, Taylor J, Seidman CE, Seidman JG & Mikawa T (2018) Dynamic Cellular Integration Drives Functional Assembly of the Heart’s Pacemaker Complex. Cell Reports 23: 2283–2291

Bressan M, Liu G & Mikawa T (2013) Early mesodermal cues assign avian cardiac pacemaker fate potential in a tertiary heart field. Science 340: 744–748

Bucchi A, Baruscotti M & Difrancesco D (2002) Current-dependent block of rabbit sino-atrial node If channels by ivabradine. Journal of General Physiology 120: 1–13

Cai CL, Liang X, Shi Y, Chu PH, Pfaff SL, Chen J & Evans S (2003) Isl1 identifies a cardiac progenitor population that proliferates prior to differentiation and contributes a majority of cells to the heart. Developmental Cell 5: 877–889

Carbon S, Douglass E, Good BM, Unni DR, Harris NL, Mungall CJ, Basu S, Chisholm RL, Dodson RJ, Hartline E, et al (2021) The Gene Ontology resource: enriching a GOld mine. Nucleic Acids Research 49: D325–D334

Chandler NJ, Greener ID, Tellez JO, Inada S, Musa H, Molenaar P, DiFrancesco D, Baruscotti M, Longhi R, Anderson RH, et al (2009) Molecular architecture of the human sinus node insights into the function of the cardiac pacemaker. Circulation 119: 1562–1575

Choudhury M, Boyett MR & Morris GM (2015) Biology of the sinus node and its disease. Arrhythmia and Electrophysiology Review 4: 28–34

Christoffels VM, Mommersteeg MTM, Trowe MO, Prall OWJ, De Gier-De Vries C, Soufan AT, Bussen M, Schuster-Gossler K, Harvey RP, Moorman AFM, et al (2006) Formation of the venous pole of the heart from an Nkx2-5-negative precursor population requires Tbx18. Circulation Research 98: 1555–1563

Churko JM, Garg P, Treutlein B, Venkatasubramanian M, Wu H, Lee J, Wessells QN, Chen SY, Chen WY, Chetal K, et al (2018) Defining human cardiac transcription factor hierarchies using integrated single-cell heterogeneity analysis. Nature Communications 9: 1–14

Cingolani E, Goldhaber JI & Marbán E (2018) Next-generation pacemakers: From small devices to biological pacemakers. Nature Reviews Cardiology 15: 139–150 doi:10.1038/nrcardio.2017.165 [PREPRINT]

Costello I, Pimeisl IM, Dräger S, Bikoff EK, Robertson EJ & Arnold SJ (2011) The T-box transcription factor Eomesodermin acts upstream of Mesp1 to specify cardiac mesoderm during mouse gastrulation. Nature Cell Biology 13: 1084–1092

Csepe TA, Zhao J, Hansen BJ, Li N, Sul L v., Lim P, Wang Y, Simonetti OP, Kilic A, Mohler PJ, et al (2016) Human sinoatrial node structure: 3D microanatomy of sinoatrial conduction pathways. Progress in Biophysics and Molecular Biology 120: 164–178

Devalla HD, Gélinas R, Aburawi EH, Beqqali A, Goyette P, Freund C, Chaix M, Tadros R, Jiang H, Le Béchec A, et al (2016) *TECRL*, a new life-threatening inherited arrhythmia gene associated with overlapping clinical features of both LQTS and CPVT. EMBO Molecular Medicine 8: 1390–1408

Devalla HD, Schwach V, Ford JW, Milnes JT, El-Haou S, Jackson C, Gkatzis K, Elliott DA, Chuva de Sousa Lopes SM, Mummery CL, et al (2015) Atrial-like cardiomyocytes from human pluripotent stem cells are a robust preclinical model for assessing atrial-selective pharmacology. EMBO Molecular Medicine 7: 394–410

Easterling M, Rossi S, Mazzella AJ & Bressan M (2021) Assembly of the Cardiac Pacemaking Complex: Electrogenic Principles of Sinoatrial Node Morphogenesis. Journal of Cardiovascular Development and Disease 8: 40

van Eif VWW, Devalla HD, Boink GJJ & Christoffels VM (2018) Transcriptional regulation of the cardiac conduction system. Nature Reviews Cardiology 15: 617–630 doi:10.1038/s41569-018-0031-y [PREPRINT]

van Eif VWW, Stefanovic S, van Duijvenboden K, Bakker M, Wakker V, de Gier-de Vries C, Zaffran S, Verkerk AO, Boukens BJ & Christoffels VM (2019) Transcriptome analysis of mouse and human sinoatrial node cells reveals a conserved genetic program. Development 146: dev173161

Espinoza-Lewis RA, Yu L, He F, Liu H, Tang R, Shi J, Sun X, Martin JF, Wang D, Yang J, et al (2009) Shox2 is essential for the differentiation of cardiac pacemaker cells by repressing Nkx2-5. Developmental Biology 327: 376–385

Farrell JA, Wang Y, Riesenfeld SJ, Shekhar K, Regev A & Schier AF (2018) Single-cell reconstruction of developmental trajectories during zebrafish embryogenesis. Science 360

Friedman CE, Nguyen Q, Lukowski SW, Helfer A, Chiu HS, Miklas J, Levy S, Suo S, Han JDJ, Osteil P, et al (2018) Single-Cell Transcriptomic Analysis of Cardiac Differentiation from Human PSCs Reveals HOPX-Dependent Cardiomyocyte Maturation. Cell Stem Cell 23: 586–598.e8

Gittenberger-De Groot AC, Mahtab EAF, Hahurij ND, Wisse LJ, Deruiter MC, Wijffels MCEF & Poelmann RE (2007) Nkx2.5-negative myocardium of the posterior heart field and its correlation with podoplanin expression in cells from the developing cardiac pacemaking and conduction system. The Anatomical Record: Advances in Integrative Anatomy and Evolutionary Biology 290: 115–122

Goodyer WR, Beyersdorf BM, Paik DT, Tian L, Li G, Buikema JW, Chirikian O, Choi S, Venkatraman S, Adams EL, et al (2019) Transcriptomic Profiling of the Developing Cardiac Conduction System at Single-Cell Resolution. Circulation Research 125: 379–397

Guadix JA, Orlova V V., Giacomelli E, Bellin M, Ribeiro MC, Mummery CL, Pérez-Pomares JM & Passier R (2017) Human Pluripotent Stem Cell Differentiation into Functional Epicardial Progenitor Cells. Stem Cell Reports 9: 1754–1764

Hafemeister C & Satija R (2019) Normalization and variance stabilization of single-cell RNA-seq data using regularized negative binomial regression. Genome Biology 20: 296

Hashimshony T, Senderovich N, Avital G, Klochendler A, de Leeuw Y, Anavy L, Gennert D, Li S, Livak KJ, Rozenblatt-Rosen O, et al (2016) CEL-Seq2: sensitive highly-multiplexed single-cell RNA-Seq. Genome Biology 17: 77

Iyer D, Gambardella L, Bernard WG, Serrano F, Mascetti VL, Pedersen RA, Talasila A & Sinha S (2015) Robust derivation of epicardium and its differentiated smooth muscle cell progeny from human pluripotent stem cells. Development (Cambridge) 142: 1528–1541

Kitajima S, Takagi A, Inoue T & Saga Y (2000) MesP1 and MesP2 are essential for the development of cardiac mesoderm. Development 127: 3215–3226

Komosa ER, Wolfson DW, Bressan M, Cho HC & Ogle BM (2021) Implementing Biological Pacemakers: Design Criteria for Successful Transition From Concept to Clinic. Circulation: Arrhythmia and Electrophysiology 14

Kruithof BPT, van Wijk B, Somi S, Kruithof-de Julio M, Pérez Pomares JM, Weesie F, Wessels A, Moorman AFM & van den Hoff MJB (2006) BMP and FGF regulate the differentiation of multipotential pericardial mesoderm into the myocardial or epicardial lineage. Developmental Biology 295: 507–522

Lambright DG, Noel JP, Hamm HE & Sigler PB (1994) Structural determinants for activation of the α-subunit of a heterotrimeric G protein. Nature 369: 621–628

Li H, Li D, Wang Y, Huang Z, Xu J, Yang T, Wang L, Tang Q, Cai C-L, Huang H, et al (2019) Nkx2-5 defines a subpopulation of pacemaker cells and is essential for the physiological function of the sinoatrial node in mice. Development (Cambridge, England) 146

Liang D, Xue J, Geng L, Zhou L, Lv B, Zeng Q, Xiong K, Zhou H, Xie D, Zhang F, et al (2021) Cellular and molecular landscape of mammalian sinoatrial node revealed by single-cell RNA sequencing. Nature Communications 12: 1–15

Linscheid N, Logantha SJRJ, Poulsen PC, Zhang S, Schrölkamp M, Egerod KL, Thompson JJ, Kitmitto A, Galli G, Humphries MJ, et al (2019) Quantitative proteomics and single-nucleus transcriptomics of the sinus node elucidates the foundation of cardiac pacemaking. Nature Communications 10: 1–19

Liu X, Chen W, Li W, Li Y, Priest JR, Zhou B, Wang J & Zhou Z (2019) Single-Cell RNA-Seq of the Developing Cardiac Outflow Tract Reveals Convergent Development of the Vascular Smooth Muscle Cells. Cell Reports 28: 1346–1361.e4

Lupu IE, Redpath AN & Smart N (2020) Spatiotemporal Analysis Reveals Overlap of Key Proepicardial Markers in the Developing Murine Heart. Stem Cell Reports 14: 770–787

McInnes L, Healy J, Saul N & Großberger L (2018) UMAP: Uniform Manifold Approximation and Projection. Journal of Open Source Software 3: 861

Mikryukov AA, Mazine A, Wei B, Yang D, Miao Y, Gu M & Keller GM (2021) BMP10 Signaling Promotes the Development of Endocardial Cells from Human Pluripotent Stem Cell-Derived Cardiovascular Progenitors. Cell Stem Cell 28: 96–111.e7

Mohan RA, Boukens BJ & Christoffels VM (2018) Developmental Origin of the Cardiac Conduction System: Insight from Lineage Tracing. Pediatric Cardiology 39: 1107–1114 doi:10.1007/s00246-018-1906-8 [PREPRINT]

Mommersteeg MTM, Domínguez JN, Wiese C, Norden J, de Gier-De Vries C, Burch JBE, Kispert A, Brown NA, Moorman AFM & Christoffels VM (2010) The sinus venosus progenitors separate and diversify from the first and second heart fields early in development. Cardiovascular Research 87: 92–101

Mommersteeg MTM, Hoogaars WMH, Prall OWJ, De Gier-De Vries C, Wiese C, Clout DEW, Papaioannou VE, Brown NA, Harvey RP, Moorman AFM, et al (2007) Molecular pathway for the localized formation of the sinoatrial node. Circulation Research 100: 354–362

Muraro MJ, Dharmadhikari G, Grün D, Groen N, Dielen T, Jansen E, van Gurp L, Engelse MA, Carlotti F, de Koning EJP, et al (2016) A Single-Cell Transcriptome Atlas of the Human Pancreas. Cell Systems 3: 385–394.e3

Ng ES, Davis R, Stanley EG & Elefanty AG (2008) A protocol describing the use of a recombinant protein-based, animal product-free medium (APEL) for human embryonic stem cell differentiation as spin embryoid bodies. Nature Protocols 3: 768–776

Protze SI, Liu J, Nussinovitch U, Ohana L, Backx PH, Gepstein L & Keller GM (2017) Sinoatrial node cardiomyocytes derived from human pluripotent cells function as a biological pacemaker. Nature Biotechnology 35: 56–68

Ren J, Han P, Ma X, Farah EN, Bloomekatz J, Zeng XXI, Zhang R, Swim MM, Witty AD, Knight HG, et al (2019) Canonical Wnt5b Signaling Directs Outlying Nkx2.5+ Mesoderm into Pacemaker Cardiomyocytes. Developmental Cell 50: 729–743.e5

Ruijter JM, Ramakers C, Hoogaars WMH, Karlen Y, Bakker O, van den hoff MJB & Moorman AFM (2009) Amplification efficiency: Linking baseline and bias in the analysis of quantitative PCR data. Nucleic Acids Research 37

Sahara M, Santoro F, Sohlmér J, Zhou C, Witman N, Leung CY, Mononen M, Bylund K, Gruber P & Chien KR (2019) Population and Single-Cell Analysis of Human Cardiogenesis Reveals Unique LGR5 Ventricular Progenitors in Embryonic Outflow Tract. Developmental Cell 48: 475–490.e7

Sizarov A, Ya J, de Boer BA, Lamers WH, Christoffels VM & Moorman AFM (2011) Formation of the building plan of the human heart: Morphogenesis, growth, and differentiation. Circulation 123: 1125–1135

Stefanovic S, Laforest B, Desvignes JP, Lescroart F, Argiro L, Maurel-Zaffran C, Salgado D, Plaindoux E, De Bono C, Pazur K, et al (2020) Hox-dependent coordination of mouse cardiac progenitor cell patterning and differentiation. eLife 9: 1–32

Stuart T, Butler A, Hoffman P, Hafemeister C, Papalexi E, Mauck WM, Hao Y, Stoeckius M, Smibert P & Satija R (2019) Comprehensive Integration of Single-Cell Data. Cell 177: 1888–1902.e21

Tosic J, Kim GJ, Pavlovic M, Schröder CM, Mersiowsky SL, Barg M, Hofherr A, Probst S, Köttgen M, Hein L, et al (2019) Eomes and Brachyury control pluripotency exit and germ-layer segregation by changing the chromatin state. Nature Cell Biology 21: 1518–1531

Vedantham V, Galang G, Evangelista M, Deo RC & Srivastava D (2015) RNA sequencing of mouse sinoatrial node reveals an upstream regulatory role for Islet-1 in cardiac pacemaker cells. Circulation Research 116: 797–803

Verkerk AO, Wilders R, Van Borren MMGJ, Peters RJG, Broekhuis E, Lam K, Coronel R, De Bakker JMT & Tan HL (2007) Pacemaker current (If) in the human sinoatrial node. European Heart Journal 28: 2472–2478

Vicente-Steijn R, Kolditz DP, Mahtab EAF, Askar SFA, Bax NAM, Van Der Graaf LM, Wisse LJ, Passier R, Pijnappels DA, Schalij MJ, et al (2010) Electrical activation of sinus venosus myocardium and expression patterns of RhoA and Isl-1 in the chick embryo. Journal of Cardiovascular Electrophysiology 21: 1284–1292

Wiese C, Grieskamp T, Airik R, Mommersteeg MTM, Gardiwal A, de Gier-De Vries C, Schuster-Gossler K, Moorman AFM, Kispert A & Christoffels VM (2009) Formation of the sinus node head and differentiation of sinus node myocardium are independently regulated by Tbx18 and Tbx3. Circulation Research 104: 388–397

Wiesinger A, Boink GJJ, Christoffels VM & Devalla HD (2021) Retinoic acid signaling in heart development: Application in the differentiation of cardiovascular lineages from human pluripotent stem cells. Stem Cell Reports 16: 2589–2606

van Wijk B, van den Berg G, Abu-Issa R, Barnett P, van der Velden S, Schmidt M, Ruijter JM, Kirby ML, Moorman AFM & van den Hoff MJB (2009) Epicardium and Myocardium Separate From a Common Precursor Pool by Crosstalk Between Bone Morphogenetic Protein– and Fibroblast Growth Factor–Signaling Pathways. Circulation Research 105: 431–441

Witty AD, Mihic A, Tam RY, Fisher SA, Mikryukov A, Shoichet MS, Li RK, Kattman SJ & Keller G (2014) Generation of the epicardial lineage from human pluripotent stem cells. Nature Biotechnology 32: 1026–1035

Yang J, Huang J, Maity B, Gao Z, Lorca RA, Gudmundsson H, Li J, Stewart A, Swaminathan PD, Ibeawuchi SR, et al (2010) RGS6, a modulator of parasympathetic activation in heart. Circulation Research 107: 1345–1349

Zhao J, Cao H, Tian L, Huo W, Zhai K, Wang P, Ji G & Ma Y (2017) Efficient Differentiation of TBX18+/WT1+ Epicardial-Like Cells from Human Pluripotent Stem Cells Using Small Molecular Compounds. Stem cells and development 26: 528–540

